# Noncanonical Notch signals have opposing roles during cardiac development

**DOI:** 10.1101/2021.08.09.455692

**Authors:** Matthew Miyamoto, Peter Andersen, Edrick Sulistio, Xihe Liu, Sean Murphy, Suraj Kannan, Lucy Nam, William Miyamoto, Emmanouil Tampakakis, Narutoshi Hibino, Hideki Uosaki, Chulan Kwon

## Abstract

The Notch pathway is an ancient intercellular signaling system with crucial roles in numerous cell-fate decision processes across species. While the canonical pathway is activated by ligand-induced cleavage and nuclear localization of membrane-bound Notch, Notch can also exert its activity in a ligand/transcription-independent fashion, which is conserved in Drosophila, Xenopus, and mammals. However, the noncanonical role remains poorly understood in in vivo processes. Here we show that increased levels of the Notch intracellular domain (NICD) in the early mesoderm inhibit heart development, potentially through impaired induction of the second heart field (SHF), independently of the transcriptional effector RBP-J. Similarly, inhibiting Notch cleavage, shown to increase noncanonical Notch activity, suppressed SHF induction in embryonic stem cell (ESC)-derived mesodermal cells. In contrast, NICD overexpression in late cardiac progenitor cells lacking RBP-J resulted in an increase in heart size. Our study suggests that noncanonical Notch signaling has stagespecific roles during cardiac development.

---

Notch is an evolutionarily conserved signaling pathway responsible for various cell-fate decisions throughout development. Notch signaling has been shown to be involved in several aspects of heart development [1,2], including development of cardiac progenitor cells (CPCs) and cardiomyocytes (CMs) [3–5], development of the cardiac conduction system [6] and ventricular myocyte differentiation [7], whereas dysregulation of Notch signaling is associated with various congenital heart abnormalities [8–11], the most common type of birth defects.

The canonical Notch signaling cascade begins with the binding of a Delta/Serrate/LAG-2 (DSL) family extracellular ligand to a Notch Extracellular Domain, leading to the cleavage of the Notch Intracellular Domain (NICD), which translocates to the nucleus where it binds the transcription factor RBP-j to initiate gene expression [12–15]. While NICD has generally been thought to be biologically inactive outside of this canonical signaling cascade, several studies have demonstrated how it is involved in biological processes in multiple systems independent of ligand activation and/or RBP-j, suggesting noncanonical roles of Notch [16–18]. Consistent with this, we and others recently described how membrane-bound and cytoplasmic NICD post-translationally regulates the active form of β-catenin independent of RBP-j [19–22], suggesting that Notch may regulate multiple processes of early development by regulating Wnt signaling. Wnt signaling plays a key inductive role in primitive streak and cardiac mesoderm formation [23–25], and later a regulatory role in CPC development [26–31]. These interactions can be recapitulated in vitro, where stage-specific manipulation of Wnt activity aids in directed differentiation of pluripotent stem cells to cardiomyocytes [31]. However, the role of noncanonical Notch signaling in heart development remains unknown. In this present study, we demonstrate that noncanonical Notch signals inhibit heart development in early development but increase heart size at later developmental stages, suggesting its distinct and biphasic roles during cardiac development.

## Results

### Increased levels of Notch Intracellular Domain decreases heart size independent of RBP-j

To investigate the role of Notch signaling during early cardiac development, we activated NICD in precardiac mesoderm by crossing Mesp1-Cre mice with mice harboring a loxP flanked stop codon in front of NICD (NICDOE). The activation resulted in the absence of a linear heart tube at E8.5 (Fig. 1A), leading to embryonic lethality at E9.0. The mutant embryos had a cardiac crescent-like structure at E8.5, normally present transiently from E7.5–8.0, but failed to progress to form a heart tube (Fig. 1A). Additionally, the transverse sectioning and staining with the CM marker Nkx2.5 showed a thickening of the cardiac crescent compared to control embryos (Fig. 1A). Since NICD can affect cells without the obligatory canonical Notch signaling transcription factor RBP-j [19–22], we asked whether the observed cardiac phenotype was a result of canonical or noncanonical Notch signaling. To do this, we simultaneously deleted RBP-j. Surprisingly, the resulting embryos also showed the cardiac crescent at E8.5 with a reduced heart size, phenocopying NICDOE embryos with RBP-j (Fig. 1A). This suggests that the observed phenotype results from the biological activity of NICD outside of the canonical Notch signaling pathway. The phenotypic similarity was further confirmed by Nkx2.5 staining (Fig. 1A). The heart tube was formed normally in RBP-j knockout embryos (Fig. 1A), indicating the NICDOE phenotype was not a result of RBP-j deletion. These findings suggest that Notch can suppress heart tube formation in a noncanonical manner.

**Fig. 1.**
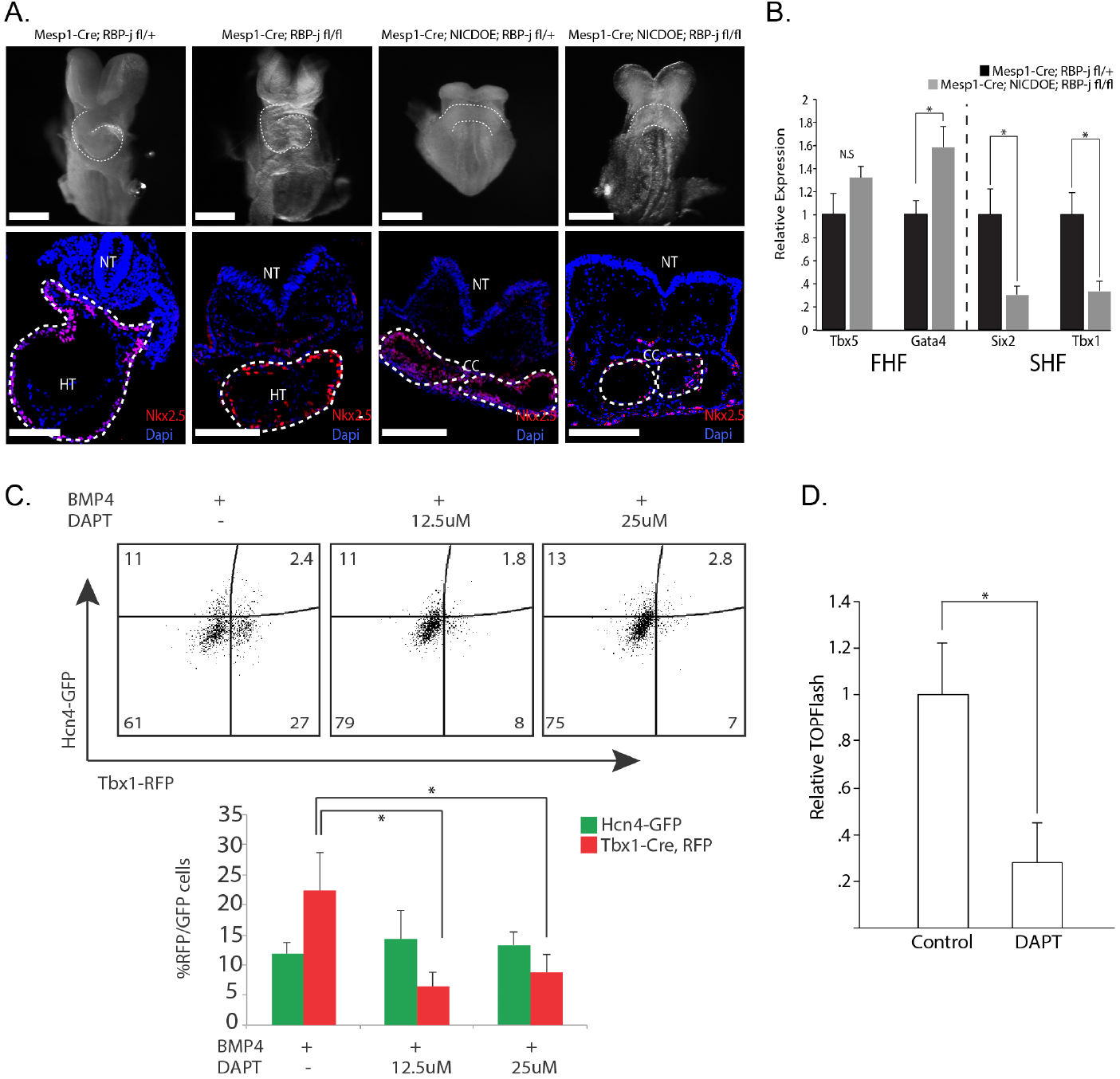
Noncanonical Notch inhibits heart development in Mesp1+ CPCs. **A.** E8.5 embryos whole mount and transverse sectioning showing decreased heart size in NICDOE embryos, independent of RBP-j. Cardiac crescent (CC) develops normally, but looping heart (HT) with visible left ventricle (LV) and outflow tract (OT) are not present in mutant embryos (n=10). White scale bars represent 300μm. **B.** Relative gene expression of FHF/SHF genes from embryos dissected at E8.0. FHF marker *Tbx5* shows no significant difference and FHF marker *Gata4* shows a slight upregulation in NICDOE embryos. SHF markers Six2 and Tbx1 both show significant downregulation in NICDOE embryos (n=9). **C.** Representative flow cytometric analysis plots quantifying GFP+/RFP+ cells after DAPT treatment. Percentage of RFP+ (SHF) cells significantly decreases with DAPT treatment. D. DAPT treatment in CPCs during induction leads to decrease in Wnt activity. **D.** DAPT treatment in CPCs during induction leads to decrease in Wnt activity

### Noncanonical Notch signaling inhibits SHF formation

Next, we sought to determine how NICD activation affects precardiac mesoderm. While NICDOE embryos formed the cardiac crescent, referred to as the first heart field (FHF), it is unclear if NICDOE affects the second heart field (SHF) that gives rise to the outflow tract and right ventricle. To test this, we examined expression of FHF/SHF genes in NICDOE embryos at E8.0 in cells of the Mesp1 lineage, specifically. Interestingly, the FHF gene *Tbx5* was not affected; though, *Gata4* expression was slightly upregulated (Fig. 1B). However, the SHF genes *Tbx1* and *Six2* were significantly downregulated (Fig. 1B). This result suggests that noncanonical Notch signals negatively regulate SHF genes.

To determine if the noncanonical signals affect heart field induction, we utilized our recently developed precardiac organoid system which harbors green and red fluorescent protein under the control of the FHF gene *Hcn4* and the SHF gene *Tbx1*, respectively [33,34]. Cardiac differentiation was done as described [32]. We previously reported that noncanonical NICDOE phenotype is recapitulated by increasing the levels of membrane-bound Notch, which can be done by blocking Notch cleavage with the small molecule N-[(3,5-Difluorophenyl)acetyl]-L-alanyl-2-phenyl]glycine-1,1-dimethylethyl ester (DAPT) [21]. Thus, we treated ESC-derived precardiac organoids with DAPT and analyzed the formation of the FHF and SHF through fluorescence flow cytometric analysis (Fig. 1C). Interestingly, we saw a significant decrease in SHF CPC formation in DAPT treated precardiac organoids when compared to controls (Fig. 1C). These results support our in vivo findings and suggest that noncanonical Notch activity inhibits SHF induction.

Previous studies have reported several factors regulating the two heart fields. While both FHF and SHF cells are regulated by FGFs [35,36] and BMPs [37], Wnt/β-catenin signaling was shown to selectively regulate SHF cells [26,38]. Given that noncanonical Notch negatively regulates Wnt/β-catenin signaling in a conserved fashion [16], we hypothesized that the decrease in SHF induction could be a result of a decrease in Wnt signaling. To test this, we treated ESC-derived mesodermal cells with DAPT during the cardiac induction period and performed a dual-luciferase TOPflash reporter assay to measure Wnt activity. The treatment greatly reduced Wnt/β-catenin activity (Fig. 1D), correlating with the decrease in SHF cells. These data suggest that noncanonical Notch signaling may suppress SHF formation via inhibition of Wnt/β-catenin signaling, and the suppression may be associated with the decreased number of cardiomyocytes observed in vivo.

### Activation of noncanonical Notch in Nkx2.5+ CPCs leads to an increase in heart size

Wnt/β-catenin signaling was shown to have a biphasic role in cardiac development, promoting cardiac lineage commitment in early mesoderm while inhibiting cardiogenesis in later development [29]. Given the conserved interaction between noncanonical Notch and Wnt/β-catenin signaling, we tested whether noncanonical Notch signaling also plays a biphasic role during cardiac development. To do this, we activated NICD in late CPCs with a Nkx2.5-Cre driver [39]. Surprisingly, in contrast to the early activation of NICD in precardiac mesoderm, the late activation resulted in a dramatic increase in heart size at E9.5, and was similarly enlarged in NICDOE embryos with simultaneous knockout of RBP-j (Fig. 2A). Further examination of mutant embryos showed an increase in size of the left ventricle, a FHF-derived chamber. Transverse sections stained for the FHF marker TBX5 showed an expansion of the left ventricle in NICD activated embryos (Fig. 2B), supporting this notion. We next looked at the functional capability of control and mutant embryos to circulate through embryo ink injection experiments. Interestingly, we observed an increased level of vascular defects in Nkx2.5-Cre NICDOE embryos in both RBP-j control and knockout conditions (Fig. 2C), suggesting defective vasculature formation or cardiomyocyte contractility in NICDOE embryos. Taken together, these results support that NICD can play a stage-specific role in CPCs during cardiac development.

**Fig. 2.**
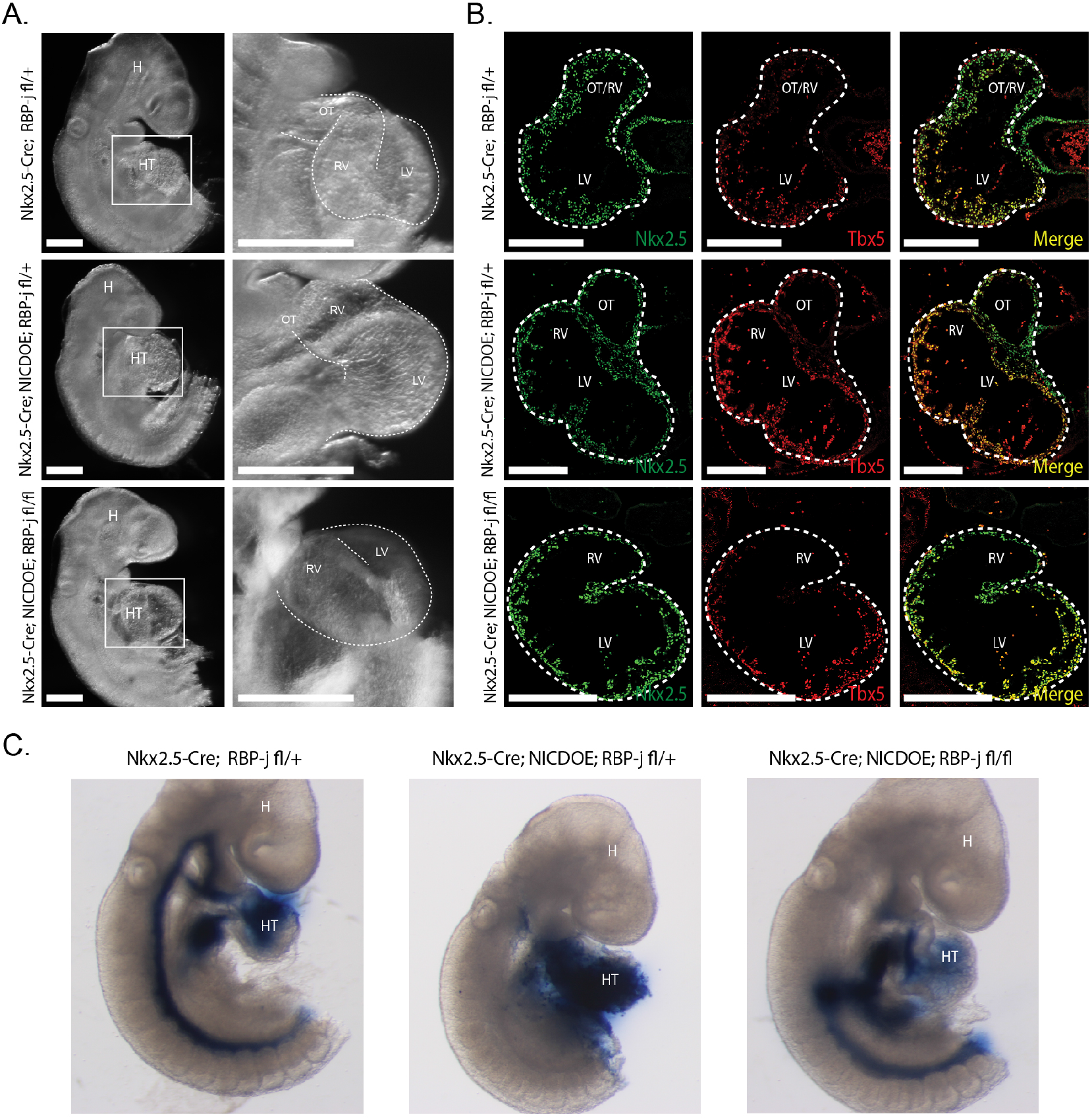
Noncanonical Notch increases heart size in Nkx2.5 CPCs. **A.** NICDOE embryos show increased heart size independent of RBP-j. Of note, LV is enlarged in NICDOE embryos compared to control (n=8/11;n=6/9). White scale bars represent 300μm. **B.** Tbx5 staining reveals an expansion of LV in transverse sections of NICDOE embryos. White scale bars represent 300μm. **C.** Ink injection experiments show disrupted circulation in NICDOE embryos independent of RBP-j (n=10/12).

## Discussion

This study provides in vivo and in vitro evidence for distinct roles of a noncanonical Notch signaling pathway. Our findings suggest that noncanonical Notch signaling plays a biphasic role in CPCs through inhibition of the second heart field in early development and promotion of cardiomyocyte proliferation in later development. Given the involvement of Notch signaling in numerous developmental and disease processes, it would be of great importance to revisit its noncanonical roles and to determine its context-dependent interactions with Wnt/β-catenin signaling.

## Methods

### Mouse work

Mesp1-Cre; NICD and Nkx2.5-Cre; NICD embryos were generated by crossing Mesp1/Nkx2.5-Cre mice [40,39] with mice harboring a NICD sequence downstream of a stop sequence flanked by loxP sites in the Rosa locus [41]. Simultaneous RBP-j knockout was achieved by crossing with mice harboring RBP-j sequence flanked by loxP sites [42]. Embryos were harvested from E8.0-E9.5 and were dissected using forceps under a Zeiss Discovery V8 microscope. Embryos intended for immunostaining were fixed in 4% PFA for 1 hour, maintained in 30% sucrose, embedded in OCT and sectioned. Immunostainings were completed using NKX2.5 antibody (Santa Cruz), TBX5 antibody, and DAPI (Life Technologies). Stained sections were imaged using a Keyence BZ-X700 microscope and post-imaging processing completed in Adobe Photoshop. Embryos intended for RT-qPCR analysis were harvested and placed in Trizol for RNA isolation. Standard RNA isolation protocols were completed. cDNA was created using the high-capacity cDNA RT kit. qPCR was completed using Sybr Select qPCR mix with primers indicated and gene expression was normalized to GAPDH. Dissected yolk sac was used for genotyping embryos.

### Cell Culture Maintenance and Differentiation

Mouse ESCs were maintained and differentiated as described [33,34]. Mouse ESCs were maintained on gelatin-coated dishes in 2i inhibitor containing: Glasgow minimum essential medium supplemented with 10% fetal bovine serum, 3 μM Chir99021 and 1 μM PD98059, 1000 U/ml ESGRO, Glutamax, Sodium Pyruvate, MEM non-essential amino acids. For differentiation of mouse embryonic stem cells, cells were plated in 3:1 IMDM/Ham’s F12 with B27, N2, Pen/Strep, Glutamax, BSA, L-ascorbic acid, and MTG for embryoid bodies (EBs) formation. After 50 hours, EBs were induced for 46 hours with BMP4 and Activin A. DAPT (CAS 208255-80-5 - Calbiochem) was administered at a concentration of 12.5 μM. Flow cytometrical analysis was carried out using a SH800 Cell sorter (Sony Biotechnologies). Luciferase assays consisted of transfecting cells in single cells suspensions with TOPFlash and Renilla constructs, and were analyzed as described [32].

### Statistical analyses

The two-tailed Student t-test, type II, was used for data analyses. P<0.05 was considered significant.

## Acknowledgements

The authors thank the Kwon laboratory members for helpful discussion. This work was supported by grants from NIH (R01HL111198, R01HD086026), MSCRF (2015-MSCRFI-1622), AHA (15GRNT25700066, 17GRNT33670432, 18IPA34170446), and The Magic that Matters Fund.

## Author Contributions

M.M. designed and carried out this work and wrote the manuscript. P.A., E.S., X.L., S.K., S.M, L.N., W.M., E.T., and H.U. assisted in animal work. X.L., S.M. and P.A. assisted in stem cell work. P.A. performed image analysis, prepared images, and conducted flow cytometry. S.M. assisted in compiling the manuscript. P.A., N.H., H.U. designed this work. C.K. designed and supervised this work and wrote manuscript.

## Notes

### Competing Interest Statement

The authors have declared no competing interest.

